## Supplementary File for "Proteomic analysis identifies dysregulated proteins in albuminuria: a South African pilot study"

Supplementary Appendix for paper titled: Proteomic analysis identifies dysregulated proteins in albuminuria: a case-control pilot study from South Africa

Author list: Siyabonga Khoza (SK)^1^, Jaya A George (JAG)^2^, Previn Naicker (PN)^3^, Stoyan H.Stoychev (SHS)^4,5^, June Fabian (JF)^6,7,†^, Ireshyn S Govender (ISG)^3,4*†^

**†Joint senior authors**

Author details:

^1^Department of Chemical Pathology, National Health Labotatory Service and School of Pathology, University of the Witwatersrand, Faculty of Health Sciences, Johannesburg, South Africa.

^2^Wits Diagnostic Innovation Hub, University of the Witwatersrand, Johannesburg, South Africa.

^3^Future Production Chemicals, Council for Scientific and Industrial Research, Pretoria, South Africa.

^4^ReSyn BioSciences, Edenvale, South Africa.

^5^Evosep Biosystems, Odense, Denmark.

^6^Wits Donald Gordon Medical Centre, School of Clinical Medicine, Faculty of Health Sciences, University of the Witwatersrand.

^7^Medical Research Council/Wits University Rural Public Health and Health Transitions Research Unit (Agincourt), School of Public Health, Faculty of Health Sciences, University of the Witwatersrand, Johannesburg, South Africa.

| **A**  Commercial System Suitability (Hela Digest) d  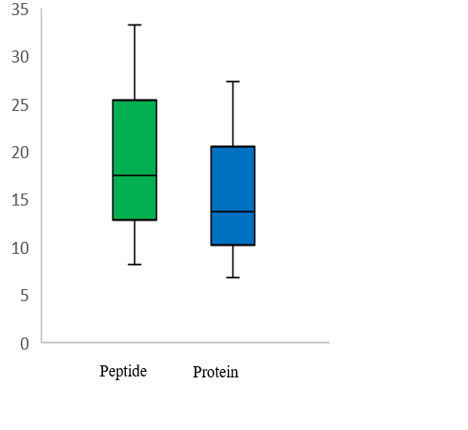  CV % | **B**  Study Specific Suitability (Urine Peptide Pool)  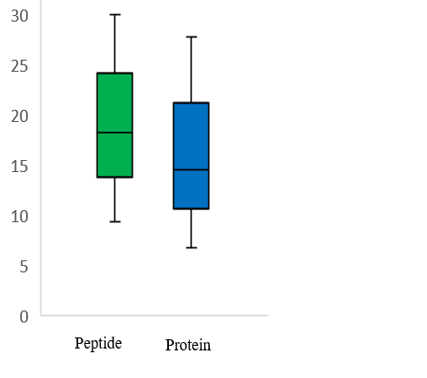  CV % |
| --- | --- |
| **C**  Commercial System Suitability (Hela Digest)  d  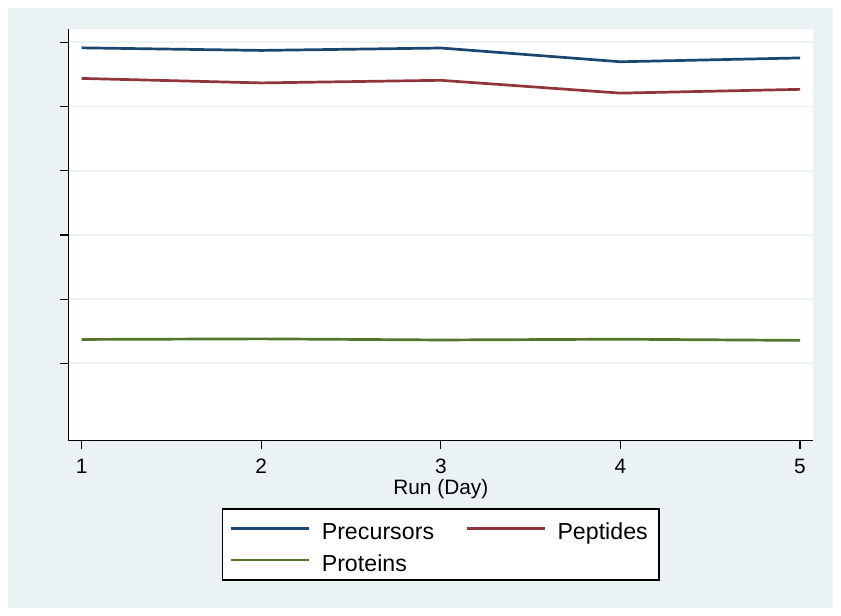 | **D**  Study Specific Suitability (Urine Peptide Pool)  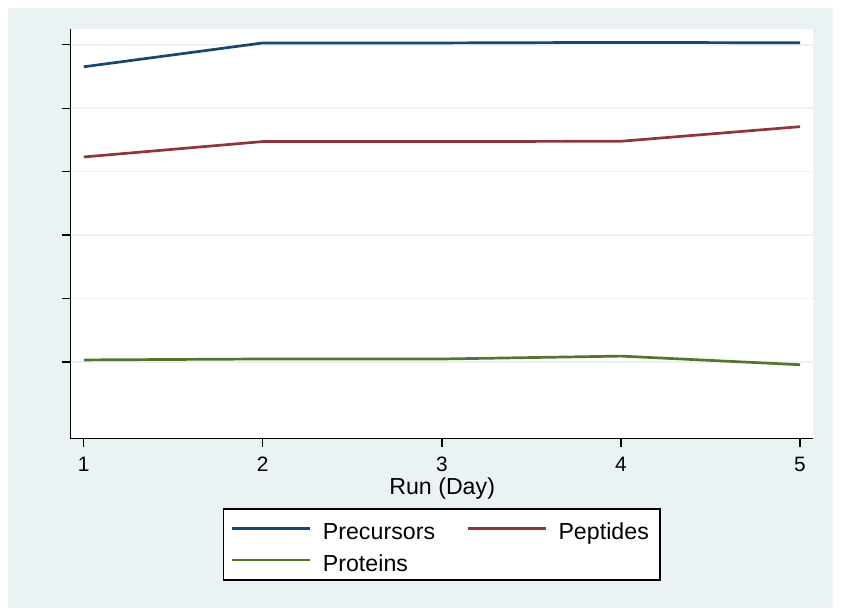 |
| **E**  Commercial System Suitability (Hela Digest) d  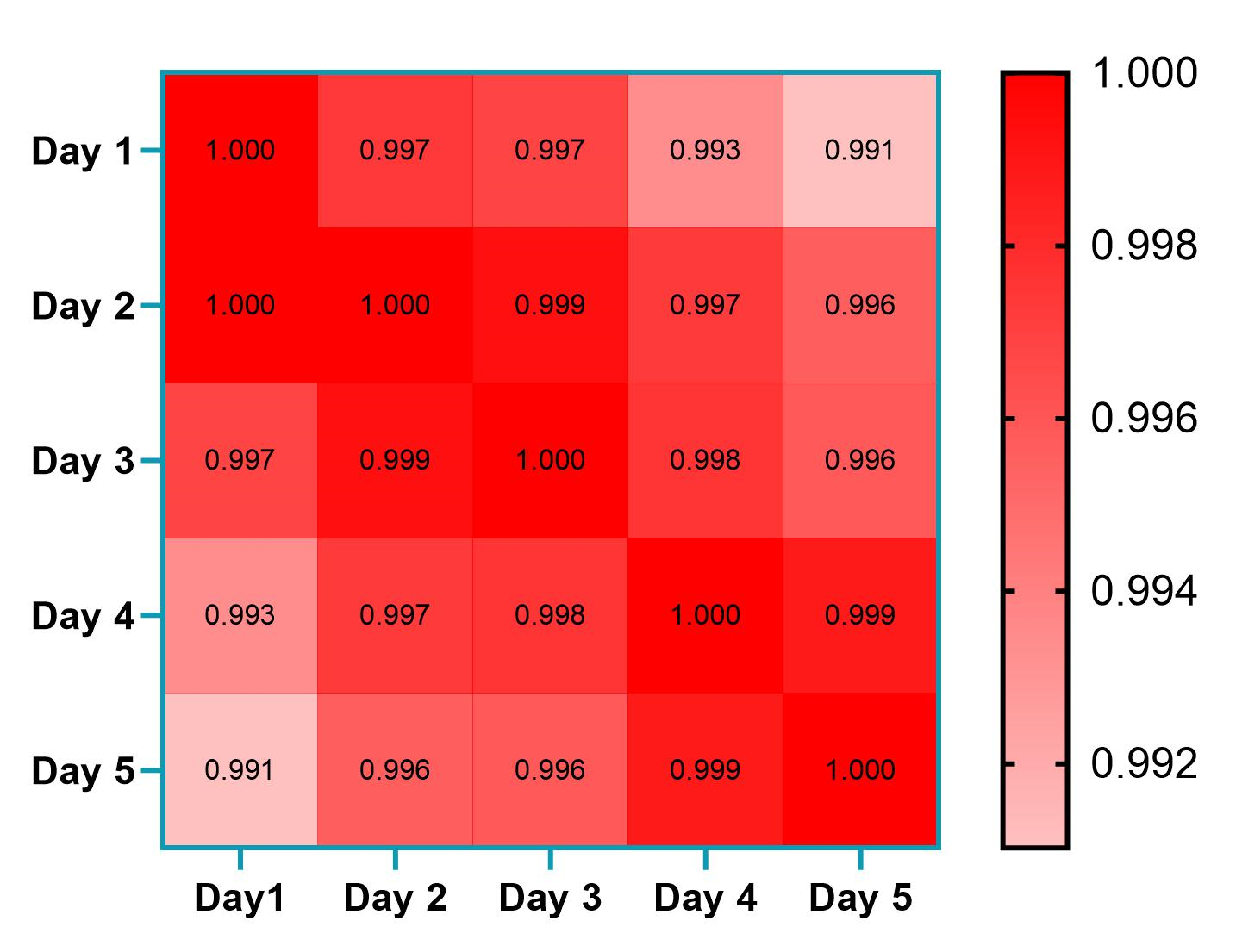 | **F**  Study Specific Suitability (Urine Peptide Pool)  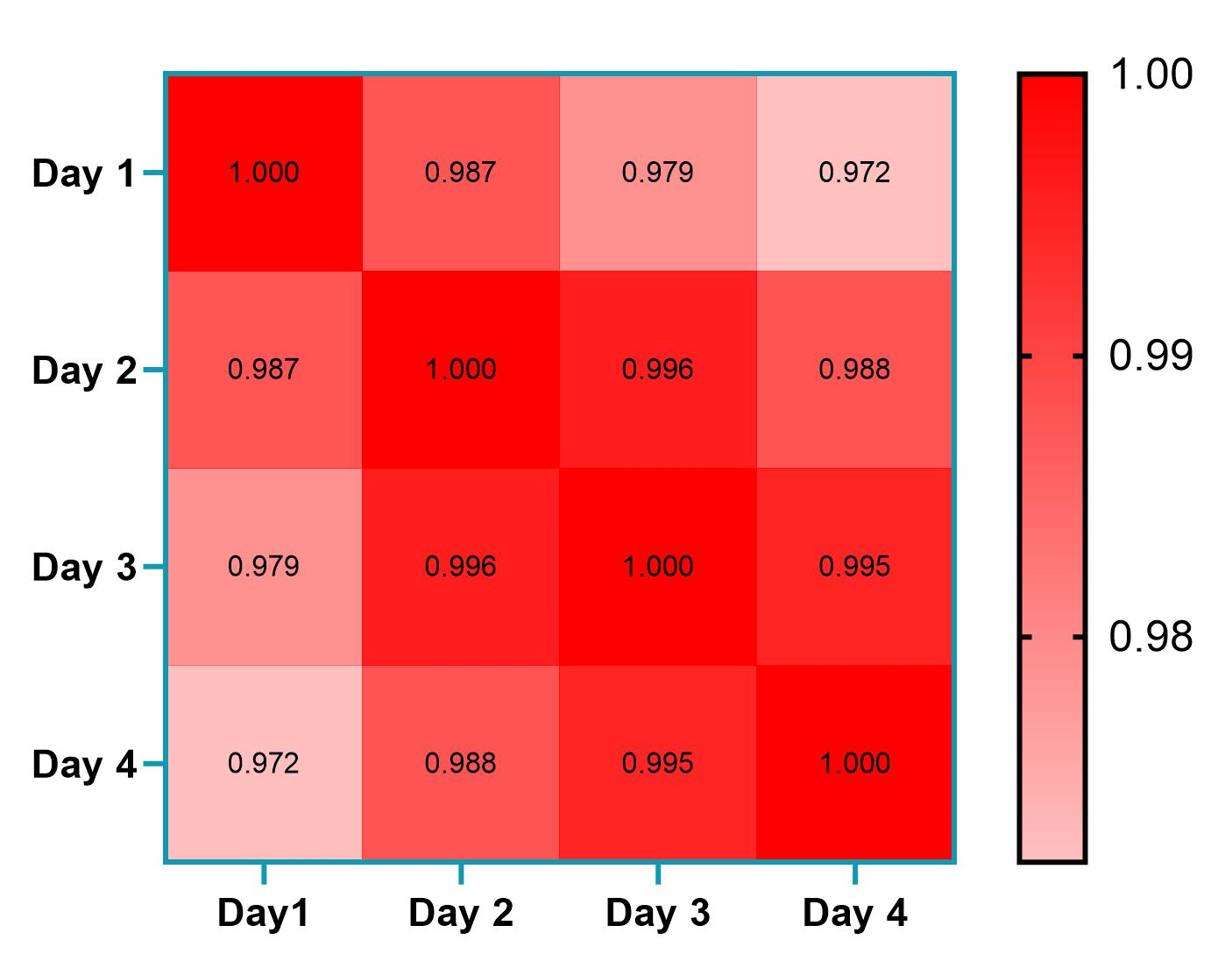 |

**Figure S1 (a-f).** Project specific system suitability and quality control (commercial Hela Digest and Urine peptide pool). CV, coefficient of variation.

| **Supplementary Table S1. Increased abundance dysregulated proteins in albuminuria sorted according to Q value** | | | | |
| --- | --- | --- | --- | --- |
| **UniProt ID** | **Protein Descriptions** | **Protein Names** | **Q value** | **Mean Log2 fold change Ratio** |
| P01009 | Alpha-1-antitrypsin (SERPINA1) | A1AT_HUMAN | 1.54E-34 | 2.904069 |
| P04217 | Alpha-1B-glycoprotein | A1BG_HUMAN | 8.35E-32 | 2.655782 |
| P02768 | Albumin | ALBU_HUMAN | 2.36E-31 | 3.103183 |
| P43652 | Afamin | AFAM_HUMAN | 1.91E-30 | 2.56667 |
| P01008 | Antithrombin-III (SERPINC1) | ANT3_HUMAN | 1.13E-24 | 2.749291 |
| P02774 | Vitamin D-binding protein | VTDB_HUMAN | 1.10E-24 | 2.359199 |
| P02144 | Myoglobin | MYG_HUMAN | 1.23E-20 | 3.631935 |
| P02647 | Apolipoprotein A-I | APOA1_HUMAN | 2.53E-12 | 3.240776 |
| P02679 | Fibrinogen gamma chain | FIBG_HUMAN | 2.96E-12 | 3.753340 |
| Q96KN2 | Beta-Ala-His dipeptidase | CNDP1_HUMAN | 6.52E-11 | 2.696401 |
| P02675 | Fibrinogen beta chain | FGB_HUMAN | 1.86E-11 | 2.494970 |
| P04439 | HLA class I histocompatibility antigen. A alpha chain | HLAA_HUMAN | 1.90E-11 | 2.647121 |
| P68871 | Hemoglobin subunit beta | HBB_HUMAN | 2.02E-11 | 3.501191 |
| O43866 | CD5 antigen-like | CD5L_HUMAN | 2.14E-11 | 4.081572 |
| P00915 | Carbonic anhydrase 1 | CAH1_HUMAN | 2.76E-10 | 2.034928 |
| P00747 | Plasminogen | PLG | 2.16E-10 | 1.499431 |
| Q9UGM5 | Fetuin-B | FETUB_HUMAN | 1.98E-10 | 4.167315 |
| P02747 | Complement C1q subcomponent subunit C | C1QC | 7.88E-10 | 2.812345 |
| P01023 | Alpha-2-macroglobulin | A2MG_HUMAN | 4.89E-10 | 3.342841 |
| P69891 | Hemoglobin subunit gamma-1 | HBG1_HUMAN | 2.71E-08 | 3.246778 |
| Q00610 | Clathrin heavy chain 1 | CLH1_HUMAN | 5.40E-09 | 3.541298 |
| A0A075B6K5 | Immunoglobulin lambda variable 3-9 | LV39_HUMAN | 7.05E-09 | 2.414491 |
| P80748 | Immunoglobulin lambda variable 3-21 | LV321_HUMAN | 4.74E-08 | 2.960301 |
| P61769 | Beta-2-microglobulin | B2MG_HUMAN | 1.54E-07 | 2.658679 |
| P07148 | Fatty acid-binding protein. liver | FABPL_HUMAN | 1.82E-07 | 2.367101 |
| P01714 | Immunoglobulin lambda variable 3-19 | IGLV3-19_HUMAN | 7.07E-07 | 2.267423 |
| P02654 | Apolipoprotein C-I | APOC1_HUMAN | 7.51E-07 | 1.815361 |
| P04003 | C4b-binding protein alpha chain | C4BPA_HUMAN | 5.85E-07 | 3.398073 |
| P02730 | Band 3 anion transport protein | B3AT_HUMAN | 5.49E-06 | 4.355263 |
| Q969H8 | Myeloid-derived growth factor | MYDGF_HUMAN | 3.33E-06 | 3.335621 |
| Q6UWP8 | Suprabasin | SBSN_HUMAN | 2.05E-06 | 3.065877 |
| P30043 | Flavin reductase (NADPH) | BLVRB_HUMAN | 1.01E-05 | 2.497917 |
| P11215 | Integrin alpha-M | ITAM_HUMAN | 4.14E-05 | 2.927536 |
| P15090 | Fatty acid-binding protein, adipocyte | FABP4_HUMAN | 2.41E-05 | 4.667801 |
| P04114 | Apolipoprotein B-100 | APOB_HUMAN | 1.54E-05 | 2.706784 |
| P24592 | Insulin-like growth factor-binding protein 6 | IBP6_HUMAN | 5.06E-05 | 4.296296 |
| P02753 | Retinol-binding protein 4 | RET4_HUMAN | 3.97E-05 | 2.637620 |
| P02042 | Hemoglobin subunit delta | HBD_HUMAN | 6.97E-05 | 3.390007 |
| P15814 | Immunoglobulin lambda-like polypeptide 1 | IGLL1 | 2.95E-05 | 2.389023 |
| P61026 | Ras-related protein Rab-10 | RAB10_HUMAN | 2.16E-05 | 2.876893 |
| P05023 | Sodium/potassium-transporting ATPase subunit alpha-1 | AT1A1_HUMAN | 1.14E-05 | 2.453589 |
| A0A087WW87 | Immunoglobulin kappa variable 2-40 | IGKV2-40_HUMAN | 8.63E-04 | 3.105648 |
| P60983 | Glia maturation factor beta | GMFB_HUMAN | 0.000043 | 3.737180 |
| P07738 | Bisphosphoglycerate mutase | PMGE_HUMAN | 0.000227 | 4.363945 |
| P50120 | Retinol-binding protein 2 | RET2_HUMAN | 0.000385 | 2.965311 |
| Q13231 | Chitotriosidase-1 | CHIT1_HUMAN | 0.000388 | 4.775099 |
| P62826 | GTP-binding nuclear protein Ran | RAN_HUMAN | 0.000572 | 2.644718 |
| P02461 | Collagen alpha-1(III) chain | CO3A1_HUMAN | 0.000870 | 2.351171 |
| P01611 | Immunoglobulin kappa variable 1-12 | KV112_HUMAN | 0.001290 | 2.514485 |
| P00568 | Adenylate kinase isoenzyme 1 | KAD1_HUMAN | 0.002011 | 2.660483 |
| P35579 | Myosin-9 | MYH9_HUMAN | 0.002226 | 4.198412 |
| P54578 | Ubiquitin carboxyl-terminal hydrolase 14 | UBP14_HUMAN | 0.003342 | 3.308528 |
| P06310 | Immunoglobulin kappa variable 2-30 | KV230_HUMAN | 0.004164 | 2.861267 |
| P06727 | Apolipoprotein A-IV | APOA4_HUMAN | 1.05E-07 | 3.525475 |
| P24666 | Low molecular weight phosphotyrosine protein phosphatase | PPAC_HUMAN | 0.004316 | 2.687916 |
| Q9NNX6 | CD209 antigen | CD209_HUMAN | 0.004914 | 2.723099 |
| P02452 | Collagen alpha-1(I) chain | COL1A1_HUMAN | 0.00661 | 2.534567 |
| P61626 | Lysozyme C | LYZ_HUMAN | 0.00548 | 2.745669 |
| P00492 | Hypoxanthine-guanine phosphoribosyltransferase | HPRT_HUMAN | 0.005473 | 2.672041 |

| **Supplementary Table S2. Decreased abundance (high in normoalbuminuria group) dysregulated proteins in albuminuria group sorted according to Q value** | | | | |
| --- | --- | --- | --- | --- |
| **UniProt ID** | **Protein Descriptions** | **Protein Name** | **Q value** | **Mean log fold change log 2 ratio** |
| P01833 | Polymeric immunoglobulin receptor | PIGR_HUMAN | 9.96E-34 | -2.555496 |
| P12109 | Collagen alpha-1(VI) chain | CO6A1_HUMAN | 1.33E-30 | -2.449243 |
| P15941 | Mucin-1 | MUC1_HUMAN | 7.87E-18 | -2.265978 |
| P39059 | Collagen alpha-1(XV) chain | COFA1_HUMAN | 1.99E-15 | -2.689387 |
| O75144 | ICOS ligand | ICOSL_HUMAN | 7.92E-15 | -2.438606 |
| P08572 | Collagen alpha-2(IV) chain | CO4A2_HUMAN | 6.32E-12 | -2.802167 |
| P07911 | Uromodulin | UMOD_HUMAN | 3.03E-11 | -2.372999 |
| P32942 | Intercellular adhesion molecule 3 | ICAM3_HUMAN | 3.28E-10 | -3.838267 |
| P10451 | Osteopontin | OSTP_HUMAN | 1.89E-09 | -2.295199 |
| P16284 | Platelet endothelial cell adhesion molecule | PECA1_HUMAN | 2.42E-09 | -2.874538 |
| Q9NZD2 | Glycolipid transfer protein | GLTP_HUMAN | 2.57E-09 | -3.375597 |
| Q7Z5N4 | Protein sidekick-1 | SDK1_HUMAN | 4.12E-09 | -2.274248 |
| P61970 | Nuclear transport factor 2 | NTF2_HUMAN | 3.82E-07 | -2.433140 |
| P22307 | Sterol carrier protein 2 | SCP2_HUMAN | 4.66E-07 | -3.897821 |
| P09488 | Glutathione S-transferase Mu 1 | GSTM1_HUMAN | 0.000222 | -2.823946 |
| P30040 | Endoplasmic reticulum resident protein 29 | ERP29_HUMAN | 0.000288 | -4.255107 |
| P32004 | Neural cell adhesion molecule L1 | L1CAM_HUMAN | 0.000411 | -3.717003 |
| Q14116 | Interleukin-18 | IL18_HUMAN | 0.002014 | -2.295583 |
| P08729 | Keratin. type II cytoskeletal 7 | K2C7_HUMAN | 0.003037 | -2.314029 |
| P12814 | Alpha-actinin-1 | ACTN1_HUMAN | 0.004721 | -3.424070 |
| Q96P63 | Serpin B12 | SPB12_HUMAN | 0.005075 | -2.591751 |

| **Supplementary Table S3. ROC curve analysis of all differentially abundant proteins** | | |
| --- | --- | --- |
| **Protein name** | **Area under curve** | **P value** |
| A1AT | 0.95137931 | 2.90E-22 |
| ALB | 0.936896552 | 1.26E-19 |
| ANT3 | 0.920689655 | 1.54E-17 |
| AFM | 0.903448276 | 1.22E-15 |
| PIGR | 0.879137931 | 1.65E-12 |
| A1BG | 0.85862069 | 1.76E-12 |
| COL6A1 | 0.842758621 | 8.41E-11 |
| MYG | 0.831724138 | 3.86E-09 |
| LV39 | 0.826724138 | 9.16E-10 |
| MUC1 | 0.813103448 | 3.04E-09 |
| ICOSLG | 0.809137931 | 7.22E-09 |
| UMOD | 0.807413793 | 2.27E-08 |
| VTDB | 0.79862069 | 1.79E-08 |
| COL15A1 | 0.778793103 | 1.78E-06 |
| CAH1 | 0.776724138 | 1.28E-07 |
| LV321 | 0.770689655 | 4.03E-07 |
| PLMN | 0.768448276 | 1.75E-06 |
| CNDP1 | 0.756724138 | 3.48E-06 |
| OSTP | 0.746896552 | 3.86E-06 |
| APOA4 | 0.741896552 | 2.48E-05 |
| PECAM1 | 0.733448276 | 8.62E-06 |
| NUTF2 | 0.731206897 | 3.71E-05 |
| APOC1 | 0.727241379 | 3.11E-05 |
| B2MG | 0.724310345 | 0.000127475 |
| FETUB | 0.719310345 | 1.55E-05 |
| KV112 | 0.714137931 | 3.90E-05 |
| SDK1 | 0.705172414 | 0.000101757 |
| CATZ | 0.701896552 | 0.002150739 |
| APOA1 | 0.688793103 | 9.53E-05 |
| KRT7 | 0.688275862 | 0.045062916 |
| B3AT | 0.68 | 0.001123533 |
| KV230 | 0.678275862 | 0.051773527 |
| CD5L | 0.677241379 | 0.001015201 |
| RAB10 | 0.649310345 | 0.007830665 |
| HLAA | 0.647931034 | 0.002391733 |
| HBB | 0.644137931 | 0.00165894 |
| PIMT | 0.64362069 | 0.002323953 |
| ERP29 | 0.642586207 | 0.013228896 |
| FIBG | 0.633965517 | 0.003786901 |
| CLH1 | 0.627586207 | 0.005692606 |
| KAD1 | 0.625689655 | 0.008093839 |
| IBP6 | 0.625689655 | 0.00808161 |
| FIBB | 0.624310345 | 0.008266953 |
| PMGE | 0.621034483 | 0.008270821 |
| HBA | 0.619827586 | 0.009591786 |
| IL18 | 0.619655172 | 0.181796271 |
| GMFB | 0.616551724 | 0.01036159 |
| HPRT | 0.612931034 | 0.024158448 |
| KV240 | 0.612586207 | 0.085686393 |
| COL4A1 | 0.609482759 | 0.015737301 |
| CHIT1 | 0.607931034 | 0.06607194 |
| HBD | 0.607241379 | 0.03756034 |
| C4BPA | 0.603793103 | 0.0194972 |
| HBG1 | 0.603793103 | 0.02539105 |
| C1QC | 0.601551724 | 0.028664713 |
| CO3A1 | 0.595344828 | 0.195909359 |
| L1CAM | 0.595 | 0.088387058 |
| FABPL | 0.593965517 | 0.039587505 |
| FABP4 | 0.593965517 | 0.033254869 |
| A2MG | 0.591724138 | 0.025853751 |
| ACTN1 | 0.587758621 | 0.327328758 |
| MYH9 | 0.587586207 | 0.131030419 |
| APOB | 0.58 | 0.031807543 |
| RET2 | 0.575862069 | 0.018305553 |
| UBP14 | 0.575344828 | 0.095410539 |
| RET4 | 0.569827586 | 0.157470673 |
| AT1A1 | 0.56637931 | 0.618572398 |
| PPAC | 0.562931034 | 0.336381777 |
| SERPINB12 | 0.561206897 | 0.055967664 |
| SCP2 | 0.560862069 | 0.402717537 |
| SBSN | 0.559827586 | 0.057042432 |
| ITAM | 0.555172414 | 0.698237266 |
| CYTC | 0.552758621 | 0.853321089 |
| BLVRB | 0.552413793 | 0.144938764 |
| GSTM1 | 0.55137931 | 0.894527925 |
| ICAM3 | 0.53862069 | 0.673757751 |
| GLTP | 0.530172414 | 0.743125431 |
| RAN | 0.522586207 | 0.727825673 |
| CD209 | 0.512068966 | 0.617768845 |
| MYDGF | 0.504482759 | 0.353317965 |

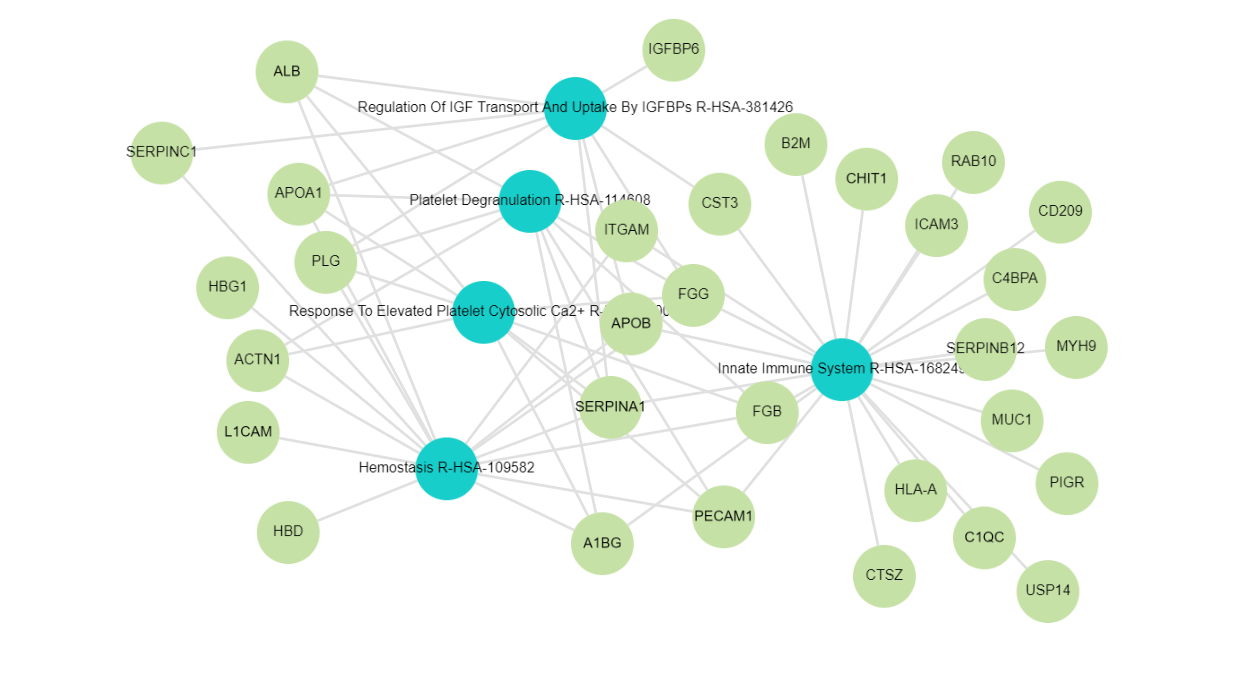

**Fig. S2** Network Analysis (Reactome library) through Enrichr-KG. Blue cycles indicates Reactome term. Green indicates genes involved.

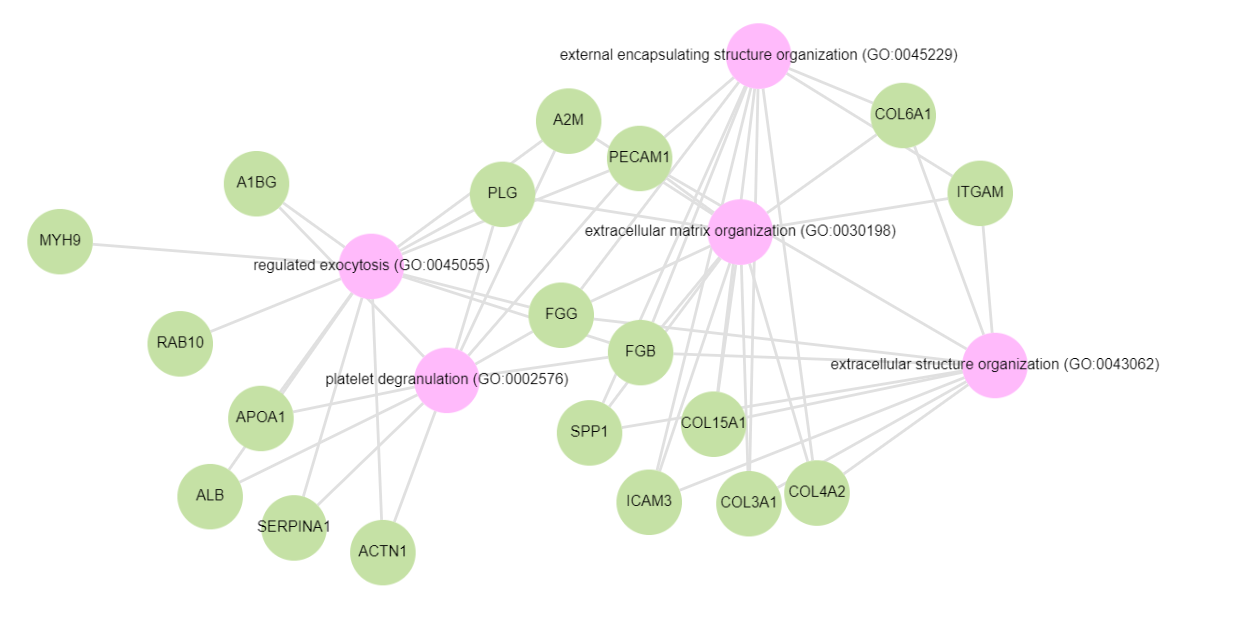
**Figure S3.** Enrichr-KG gene set enrichment analysis. GO biological-Enrich-KG: most of the proteins are involved in extracellular structure and matrix organization. Pink denote GO biological process terms. Green denotes genes.

| **Supplematary Table S4: List of genes involved in the top 5 significantly enriched biological processes.** | | | |
| --- | --- | --- | --- |
| **GO Biological Process** | **Genes** | **p-vale** | **q-value** |
| Regulated exocytosis (GO:0045055), | PECAM1, FGB, A2M, SERPINA1, RAB10, ALB, MYH9, ACTN1, APOA1, A1BG, FGG, and PLG | 6.91E-12 | 6.29E-09 |
| Platelet degranulation (GO:0002576) | APOA1, A1BG, FGG, ALB, FGB, PECAM1, ACTN1, SERPINA1, PLG, and A2M | 7.18E-11 | 3.27E-08 |
| Extracellular matrix organization (GO:0030198) | COL6A1, A2M, PLG, COL4A2, FGG, PECAM1, SPP1, COL15A1, ITGAM, FGB, COL3A1, and ICAM3 | 2.53E-09 | 7.67E-07 |
| Extracellular structure organization GO:0043062) | COL3A1, ITGAM, SPP1, ICAM3, COL6A1, COL15A1, COL4A2, PECAM1, FGG, and FGB | 1.49E-08 | 2.65E-06 |
| External encapsulating structure organization (GO:0045229) | COL6A1, COL15A1, ICAM3, ITGAM, SPP1, FGB, PECAM1, COL3A1, COL4A2, and FGG | 1.56E-08 | 2.65E-06 |
